# Thoracic crop formation is spatiotemporally coordinated with flight muscle histolysis during claustral colony foundation of a *Lasius japonicus* queen

**DOI:** 10.1101/2022.03.03.482739

**Authors:** Yuta Kurihara, Kota Ogawa, Yudai Chiba, Yoshinobu Hayashi, Satoshi Miyazaki

## Abstract

In a majority of ants, a newly mated queen independently founds a colony and claustrally raises her first brood without foraging outside the nest. During claustral independent colony foundation (ICF) in several ants, the esophagus of the founding queen expands and develops into a thoracic crop, which is then filled with a liquid substrate for larval feeding. It has been suggested that these substrates are converted from her body reserves (e.g., histolyzed flight muscles) or redistributed from a gastral crop. Here, we examined thoracic crop development in *Lasius japonicus* during claustral ICF. The foundresses claustrally fed their larvae from week 2 to 5 after ICF onset, and the first worker emerged at week 6. The development proceeded as follows: in week 0, foundress dorsal esophagus wall was pleated and thickened. Then, from week 2 to 5, the esophagus expanded dorsally toward where flight muscles had been present, following flight muscle histolysis. Gastral crop expansion followed esophagus expansion. Thus, thoracic crop formation may be spatiotemporally coordinated with flight muscle histolysis in *Lasius japonicus* queens, and similar developmental regulations might be common in other claustral ICF ants.

## INTRODUCTION

Colonial foundation is a key step in the colony life cycle of eusocial hymenopterans. Colony foundation strategies can be broadly divided into independent colony foundation (ICF), during which a foundress starts a colony alone, and dependent colony foundation (DCF), during which an existing colony is divided into two or more daughter groups (Cronin et al., 2013). Although DCF has evolved in more than 50 ant genera, ICF is a more widespread strategy and can be further divided into “non-claustral” and “claustral” ICF (Hölldobler and Wilson, 1990; Keller et al., 2014). In non-claustral ICF that is the most ancestral strategy in ants, a newly mated queen sheds her wings, excavates a new nest, and independently starts the colony (Peeters, 2020). During this founding period, the queen lays eggs, forages outside the colony, and raises her broods (Peeters and Molet, 2010). On the other hand, during claustral ICF, which is derived from non-claustral ICF only in ants (Hölldobler and Wilson, 1990; Peeters, 2020; Wheeler and Buck, 1996), a founding queen also starts the colony independently but never forages outside the nest and raises her broods using only her body reserves (Brown and Bonhoeffer, 2003). Given that foraging ants experience a high mortality risk due to predation (Nonacs and Dill, 1990; Traniello, 1989), this claustral strategy allows ant queens to avoid foraging-associated risks (Brown and Bonhoeffer, 2003; Hölldobler and Wilson, 1990). The acquisition of traits involved in claustral feeding has likely contributed to the ecological success of ants that employ such strategies. Indeed, the three subfamilies (Formicinae, Dolichoderinae, and Myrmicinae) that account for almost 80% of known ant species generally show claustral ICF (Peeters, 2020).

Claustral founding queens provide essential macronutrients such as lipids and proteins for their first larvae, and such macronutrients come from their body reserves. In claustral ICF species, young virgin queens possess enormous flight muscles (relative to their body mass) in the thorax and well-developed abdominal fat bodies (Hölldobler and Wilson, 1990; Peeters, 2020). Fat bodies are a major source of lipids and storage proteins, e.g., hexamerin (Hahn et al., 2004; Wheeler and Buck, 1996, 1995). The lipids and proteins stored in the abdomens of founding queens are depleted during claustral ICF, supporting the notion that these are transferred to the broods (Wheeler and Buck, 1996). In addition, voluminous thoraces with enormous flight muscles are a specialized traits for claustral ICF (Peeters and Ito 2001). Because queens do not fly again after their mating flight, flight muscles start to be histolyzed shortly after landing and release a large amount of protein resource (Hölldobler and Wilson, 1990; Jensen and Børgesen, 2000; Wheeler and Buck, 1996). Flight muscle histolysis of *Lasius niger* was originally described by Janet (1907) and then in greater detail by Matte and Billen (2021).

In several ICF species, the founding queen’s thoracic esophagus expands into a spindle shape and is then called a “thoracic crop” (Janet, 1907; Jensen and Børgesen, 2000; Petersen-Braun and Buschinger, 1975). This thoracic crop is thought to temporarily store the nutrients converted from flight muscles and fat bodies as liquid food for larvae (Hölldobler and Wilson, 1990). Conversely, Vander Meer et al. (1982) found that, in the fire ant *Solenopsis invicta*, the major component of the thoracic crop (mainly triacylglycerols) originated in the gastral crop. Therefore, although thoracic crop formation may serve as a key step in claustral feeding, this process is still poorly understood.

Here, we investigated the development of thoracic crop in the claustral ant *Lasius japonicus* (Formicidae: Formicinae), which is one of the most common ants in Japan and closely related with the congener *L. niger* (Maruyama et al., 2008). First, the timeline of claustral ICF, especially of claustral feeding, were examined, and then, changes in sizes, morphologies, and internal structures of the queen esophagus were examined during claustral ICF. Furthermore, it was investigated whether such changes of esophagus were associated with either flight muscle histolysis or size changes in gastral crop. Our results suggest that thoracic crop formation was spatiotemporally coordinated with flight muscle histolysis.

## MATERIALS AND METHODS

Dealate queens of *Lasius japonicus* were collected from late June to mid-July in 2018, 2019, and 2021 at the Tamagawa University campus. Each founding queen with a body length of ca. 10 mm was kept in a plastic case (36 × 36 × 14 mm, AS ONE) with fully moistened plaster. Founding colonies were maintained at approximately 25°C during experiments, and queens were not fed.

During claustral ICF, the founding queen claustrally feeds her first larvae until the eclosion of the first worker. To better understand the claustral feeding timeline, a census was performed for ten experimental colonies and the number of eggs, larvae, pupae, and workers were counted weekly until six weeks after the queen’s mating flight, when the first workers eclosed in the experimental colonies (see **RESULTS**). Because the founding queens in five of the ten experimental colonies died between weeks 5 and 6, the census at week 6 was performed only for the five remaining colonies.

To investigate the formation of thoracic crop, the queen esophagus morphology and size were examined at weeks 0–6 and also at week 8 when the first workers had eclosed in all colonies tracked. Ten to twelve founding queens were sampled from the other experimental colonies at weeks 0–6 and week 8 and then fixed in FAA fixative (formalin: acetic acid: ethanol = 6:1:16). The thoraces of the fixed queens were longitudinally dissected using a fresh razor blade and then photographed using a SZX10 stereo microscope (Olympus, Tokyo, Japan) equipped with a FLOYD digital camera (Wraymer, Osaka, Japan) and Wrayspect software (Wraymer). Because esophagus expansion occurred in the mesothorax rather than in the prothorax and metathorax (see **RESULTS**), the esophagus height in the mesothorax was measured as a proxy for esophagus size. Flight muscle histolysis was examined simultaneously.

To investigate the formation of the thoracic crop in more detail, we used the Transparentizing and Autofluorescence Scanning (TAFS) method, which enables the observation of the morphologies of decolorized and transparentized samples without staining using the autofluorescence of the samples themselves. Founding queens at weeks 0 and 3 were fixed with FAA and stored in 70% EtOH. These samples were then decolorized in a 6% hydrogen peroxide/70% EtOH solution for at least one week (Fig. S1A, B). Then, the samples were dehydrated in ascending series of isopropanol and transparentized in a BABB solution (benzyl alcohol/benzyl benzoate = 1:2) for four days (Fig. S1C). The *Lasius* queens were too large for whole-mount confocal microscopy, so they were sliced into 500 µm thick pieces using LinearSlicer PRO7 (D.S.K., Kyoto, Japan) (Fig. S1D). Sliced samples were examined with an Olympus FV3000 Confocal Laser Scanning Microscope (Olympus) using excitation lasers at wavelengths of 405, 473, and 559 nm. Fluorescence excited by lasers with different wavelengths was recorded in different channels and superimposed later. The acquired confocal images were three-dimensionally constructed and analyzed by Imaris software ver. 7.0.0 (Bitplane AG, Belfast, United Kingdom). Since autofluorescence differs among tissues, and under similar levels of excitation, the TAFS method visualizes differences in autofluorescence characteristics across tissues as differences in “color.”

To examine whether the thoracic crop contents originated in the gastral crop, the gasters (consisting of the posterior parts than third abdominal segments) of the FAA-fixed queens were dissected, and the lateral areas of the dissected crops were measured as a proxy for gastral crop size.

All statistical analyses were conducted using R ver. 4.1.0 (available at http://cran.r-project.org/).

## RESULTS

### Timeline of claustral larval feeding during *L. japonicus* colony foundation

Following their mating flights, founding queens laid 35.2 ± 14.5 eggs (mean ± SD) in the first week and then continued to lay approximately 40 eggs (38.8 ± 20.6) per week for six weeks (Fig. 1 and Fig. S2). Larvae first emerged at week 2 and then peaked in abundance at week 3 (Fig. 1). In addition, some larvae cannibalized an egg or another larva during the six weeks (Fig. S3). Pupation and eclosion were first confirmed at weeks 4 and 6, respectively (Fig. 1). These results indicate that founding queens claustrally feed their larvae at least from weeks 2 to 5, assuming that anatomical changes associated with claustral feeding should occur during or ahead of this period.

**Figure 1.**
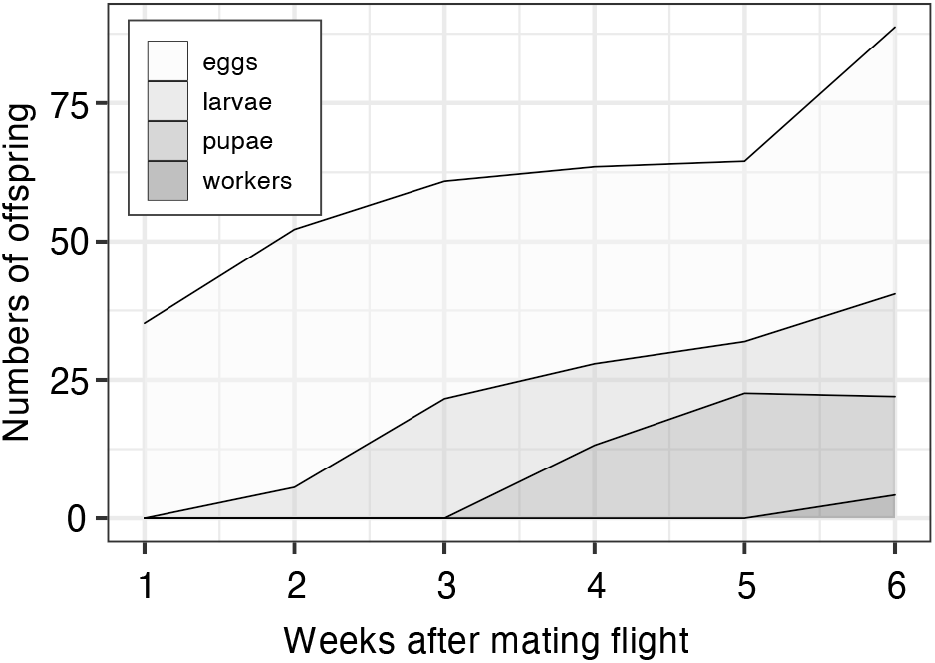
Mean offspring number produced by *Lasius japonicus* founding queens during claustral ICF. Individual plots for each developmental stage (egg, larva, pupa, and adult worker) are shown in Figure S2. Week 0 corresponds to the day of their nuptial flight.

### Thoracic crop formation parallel to flight muscle histolysis

Flight muscle degeneration began in week 1, and then, almost all muscle fibers were empty by week 3 (Fig. 2). The esophagus gradually expanded during ICF in parallel with flight muscle histolysis (Fig. 2). The esophagus after week 2 appeared larger than it did during weeks 0 and 1 (Fig. 2). Esophagus height increased significantly over time (one-way ANOVA, F = 9.287, *P* < 0.001), and at its peak height (week 5) was four times greater than its height at week 0 (Fig. 3). When the expanded esophagus was accidentally broken at weeks 3 and 5, it exuded a yellowish, oily substrate.

**Figure 2.**
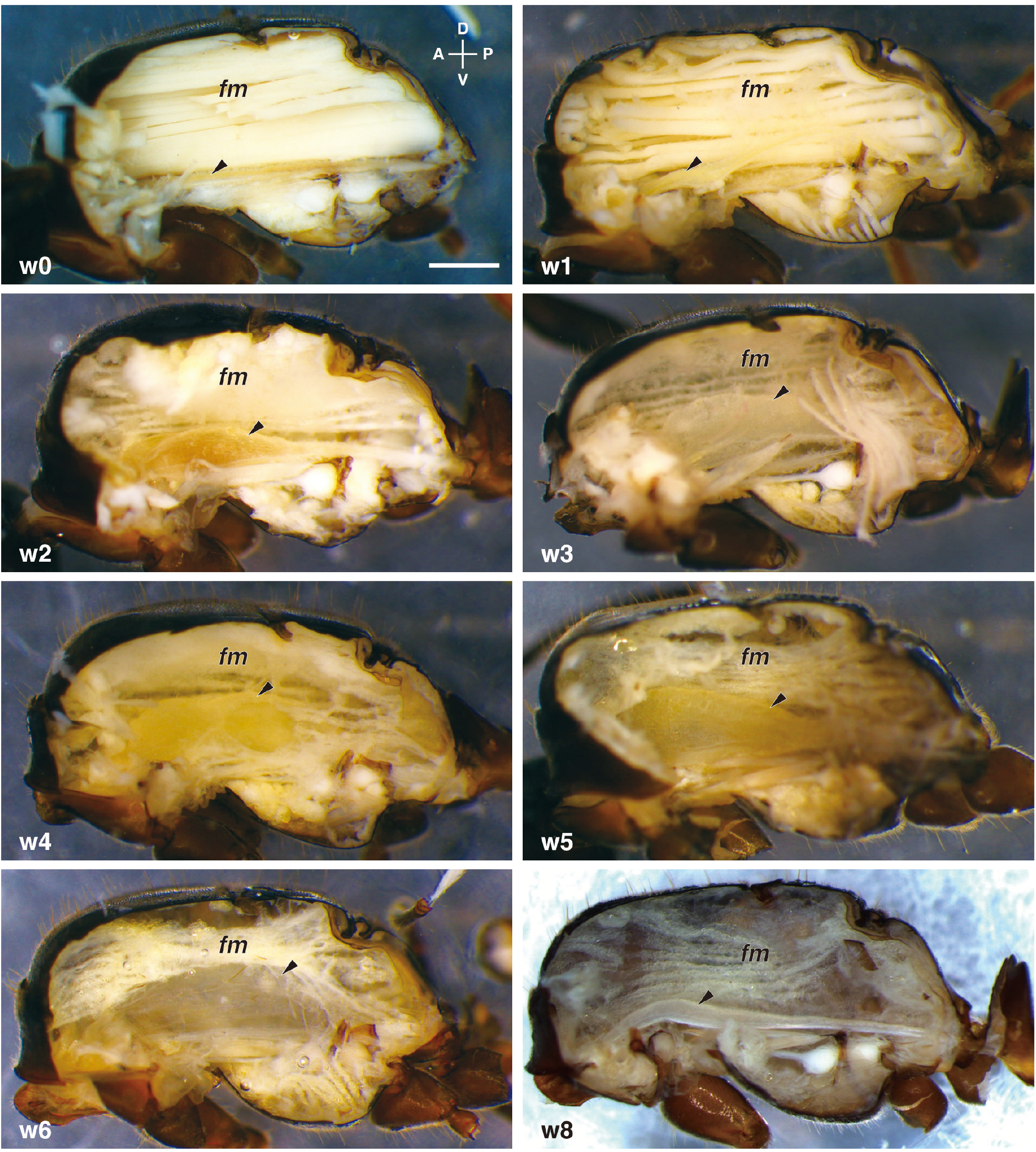
Queen esophagus and flight muscles during claustral ICF. “w0–w6” at the bottom left of each panel indicates the claustral ICF stage, i.e., 0–6 weeks after mating flight. *fm* and arrowhead indicates flight muscles and esophagus, respectively. Scale bars 500 µm.

**Figure 3.**
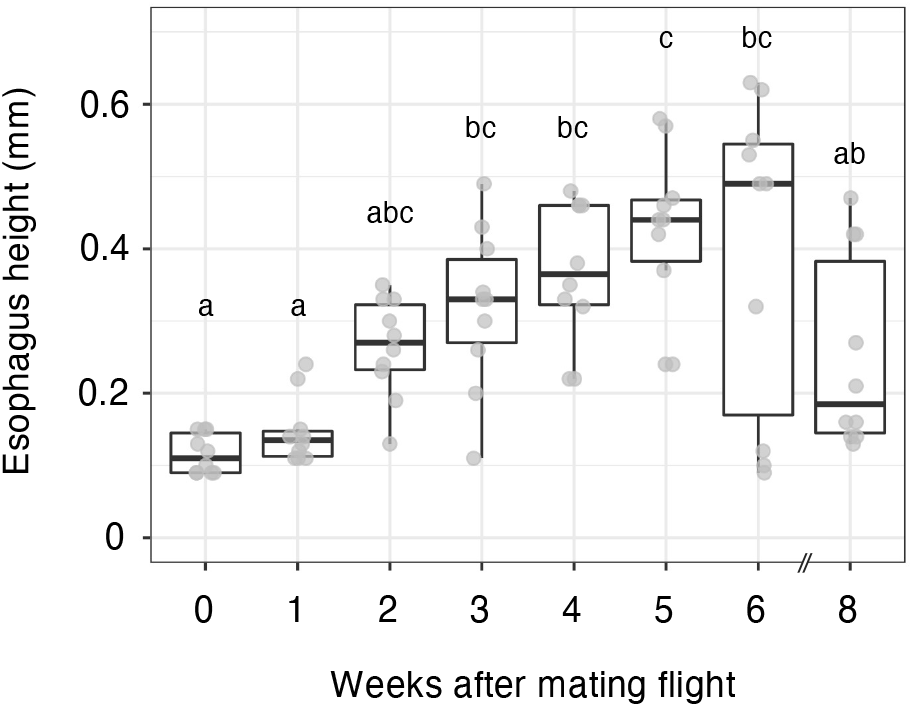
Changes in the esophagus height of a queen during claustral ICF. Each box-and-whisker plot indicates the median (horizontal line), interquartile range (box), and distance from the upper and lower quartiles times 1.5 interquartile range (whisker). All individual data shown as gray plots. Different letters indicate significant statistical differences (*P* < 0.05, Tukey–Kramer tests for post-hoc multiple comparisons).

The TAFS analyses revealed that the internal structure and size of the queen’s esophagi had changed during claustral ICF. At week 0, the thoracic esophagus was not yet distended as mentioned above (Fig. 4A), and the flight muscles had not yet decomposed, so muscle fiber structures were still recognizable (Fig. 4B). The esophagus wall was thicker and pleated throughout the thoracic esophagus (Fig. 4C), and two layers with different autofluorescence characteristics were observed (Fig. 4D–H). Of these two layers, the outer layer (topologically inside of the body) was almost uniform in thickness throughout the thoracic esophagus, whereas the thickness of the inner layer (topologically outside the body) was spatially varied. The inner layer was more thickened on the dorsal side (Fig. 4G) than on the ventral one (Fig. 4H) in the prothorax, mesothorax, and metathorax (Fig. 4D–F). In addition, the inner layer of the dorsal wall had thickened further in the mesothorax (Fig. 4E, G, and Fig. S4D) than it had in the prothorax and metathorax (Fig. 4D, F). In contrast, the esophagus wall on both the ventral and dorsal sides, where such bilayer structures could not be recognized, was thinner and less pleated in the petiole (Fig. 4I–K).

**Figure 4.**
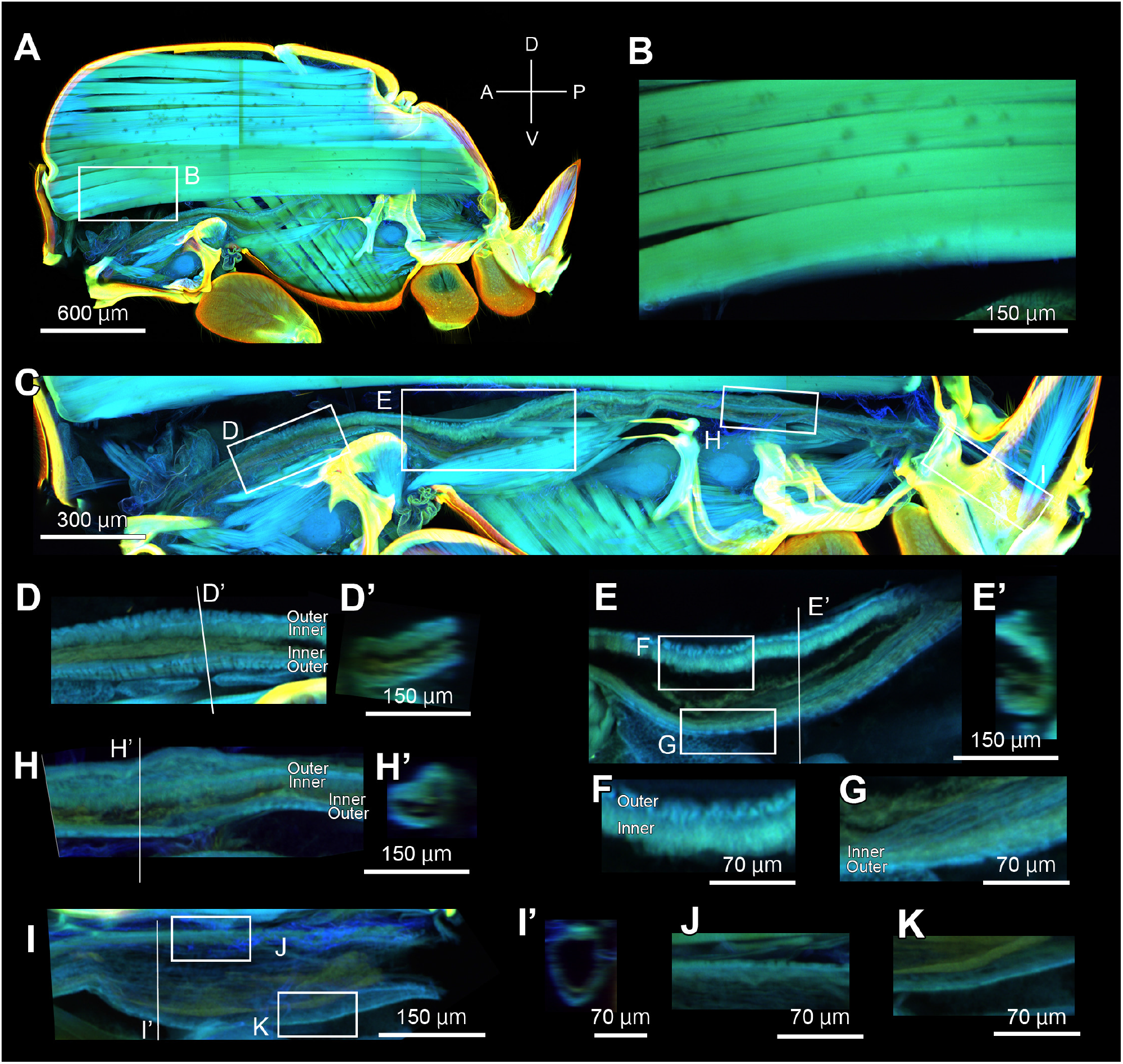
TAFS images of the thorax and petiole of a queen at week 0 in claustral ICF. Parasagittal sections of the (A) thorax and petiole, (B) flight muscle (dorsal longitudinal muscle), and (C) esophagus. Parasagittal sections of the esophagus in the (D) prothorax, (E) mesothorax ([G] dorsal and [H] ventral walls), (F) metathorax, and (I) petiole ([J] dorsal and [K] ventral walls). Thoracic esophagus consists of two layers, namely, outer (dark blue) and inner layers (light blue). Letters with single quotes indicate transverse sections marked by respective letters.

At week 3, the thoracic esophagus swelled to a spindle shape (Fig. 5A), and the flight muscles were degraded enough that no fiber structure was recognizable (Fig. 5B). The esophagus wall was thinly stretched and expanded to the dorsal side, particularly in the mesothorax (Fig. 5C and Fig. S4E), and wall thickness was almost uniform throughout the thoracic esophagus (Fig. 5D–F). The surface of the expanded esophagus showed rumpled structures extending parallel to the anterior–posterior axis (Fig. 5C), and the two-layered inner structures observed at week 0 had become unrecognizable (Fig. 5G). These results indicate that the founding queens form thoracic crops by week 2 by expanding their esophageal inner wall, particularly where it was thickened and pleated at week 0.

**Figure 5.**
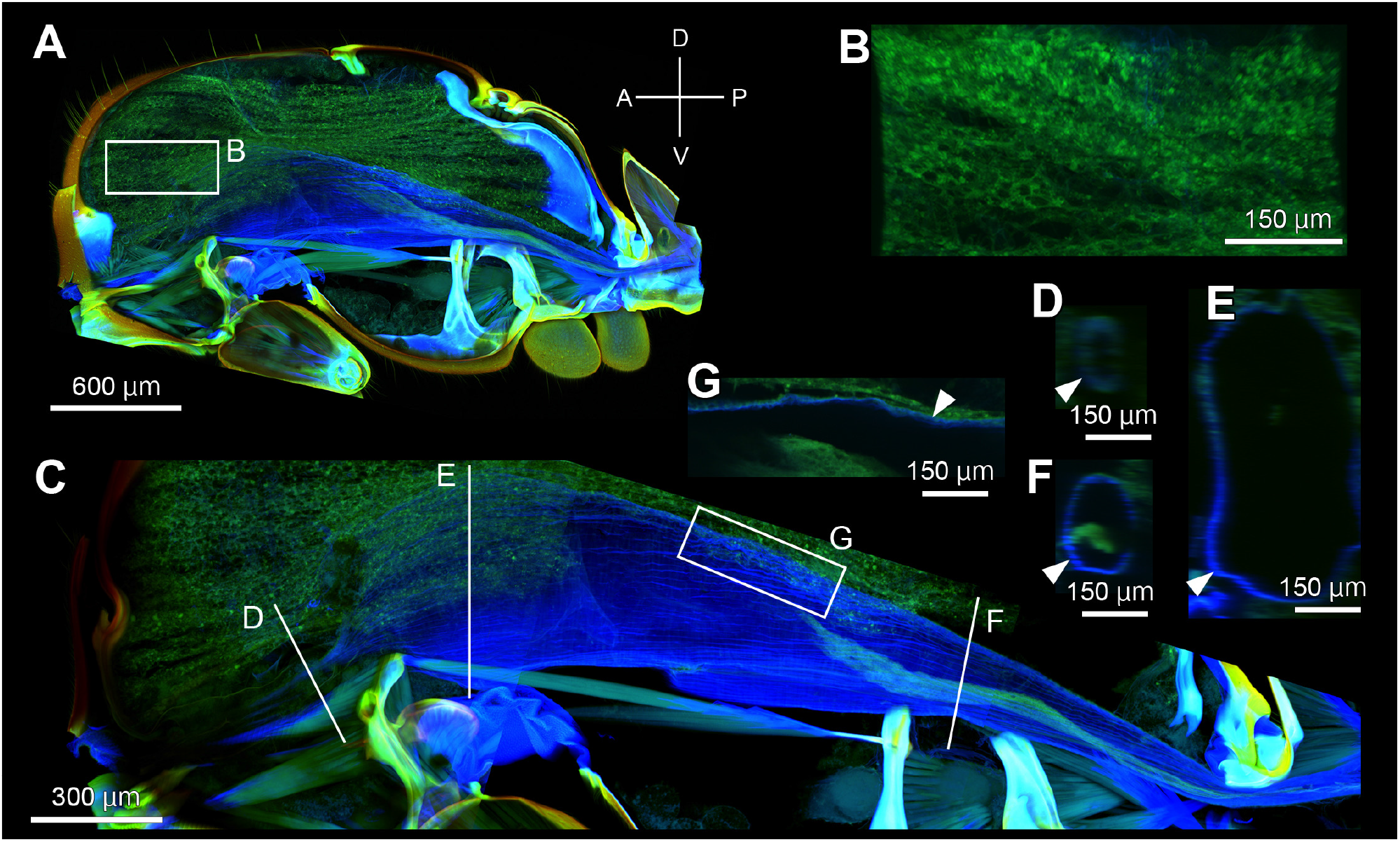
TAFS images of queen thorax and petiole at week 3 of claustral ICF. Parasagittal sections of the (A) thorax and petiole, (B) degenerating flight muscle (dorsal longitudinal muscle), and (C) expanding thoracic esophagus ([G] dorsal wall in the mesothorax). Transverse sections of the esophagus in the (D) prothorax, (E) mesothorax, and (F) metathorax. Arrowheads indicate the esophagus wall, depicted in blue.

### Changes in gastral crop size

Gastral crop size did not change for four weeks after the mating flight, but it did become significantly larger thereafter (one-way ANOVA, F = 8.448, *P* < 0.001, Fig. 6), indicating that crop size changed following esophagus expansion.

**Figure 6.**
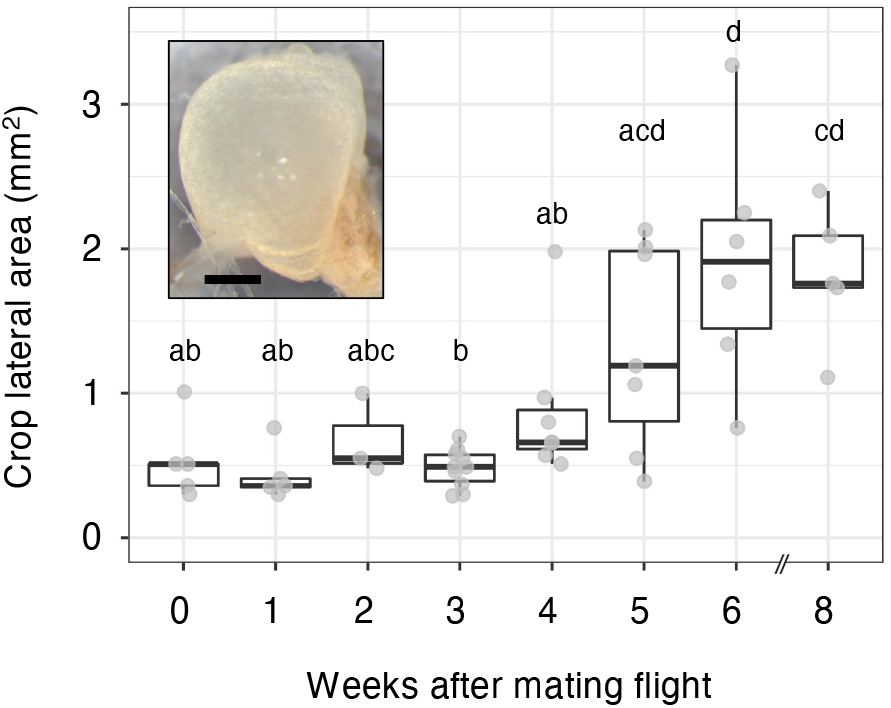
Size changes in the gastral crop of a queen during claustral ICF. The areas of the gastral crop photographed from the lateral view (inset, scale bars 300 µm) were measured as a gastral crop size. Each box-and-whisker plot indicates the median (horizontal line), interquartile range (box), and distance from the upper and lower quartiles times 1.5 interquartile range (whisker). All individual data are shown as gray plots. Different letters indicate significant statistical differences (*P* < 0.05, Tukey–Kramer tests for post-hoc multiple comparisons).

## DISCUSSION

During claustral ICF in *Lasius japonicus*, a founding queen’s flight muscles were histolyzed in the first three weeks as shown in *L. niger* (Matte and Billen, 2021), and esophageal expansion began in week 2 in parallel to the initiation of larval demand for nutrition (Fig. 7). The expansion was more evident in the esophagus’s dorsal wall than in the ventral wall, and the expanded esophagus occupied a space in the dorsal side of the thoracic cavity where flight muscle histolysis had occurred. Thus, our study shows that thoracic crop formation is spatiotemporally coordinated with flight muscle histolysis in the claustral founding queen of *L. japonicus*. In addition, the inner layer of the dorsal esophagus wall at week 0 was thicker and more pleated than that of the ventral wall, in the mesothorax where the thoracic crop was formed after week 2. The rapid expansion of the esophagus within the first three weeks of claustral ICF may be facilitated by stretching these thickened and pleated esophagus walls. The expansion of the dorsal esophagus wall in the claustral founding queen has been reported in five species from two subfamilies (Myrmicinae and Formicinae) (Jensen and Børgesen, 2000; Petersen-Braun and Buschinger, 1975). In addition, pleats in the dorsal esophagus wall were reported in newly eclosed queens of the Myrmicinae ant *Monomorium pharaonis* (Petersen-Braun and Buschinger, 1975). Therefore, the pleating, thickening, and expansion of the dorsal esophagus wall are likely adaptations that enable thoracic crop formation in spatiotemporal coordination with flight muscle histolysis, which would be a common process in claustral ant species.

**Figure 7.**
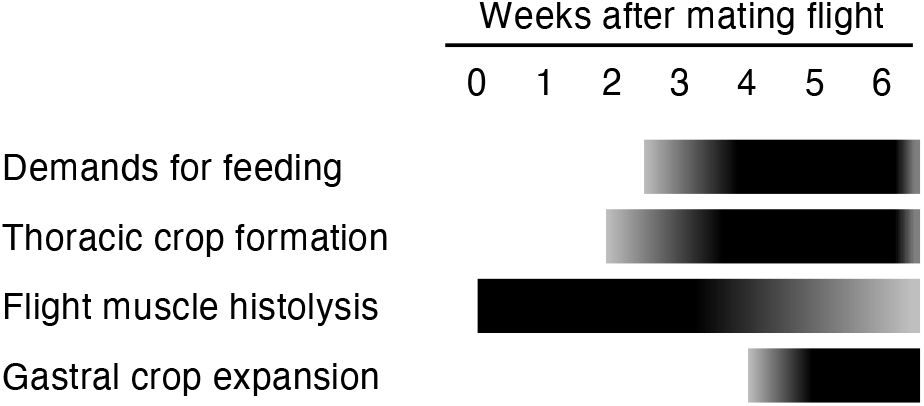
Summarized timeline of thoracic crop formation, flight muscle histolysis, and gastral crop expansion, as well as changes on larval demands for feeding, during claustral ICF in *Lasius japonicus*.

As reported in other ants, the thoracic crop of *L. japonicus* was filled with yellowish and oily substrates during the claustral feeding period (Petersen-Braun and Buschinger, 1975), which would provide nutrition for larvae. Thoracic crop formation occurred prior to gastral crop expansion (Fig. 7). In contrast to the previous study showing that lipids (mainly triacylglycerols) stored in a gastral crop flowed into the thoracic crop in fire ant queens (Vander Meer et al., 1982), nutritive resources within the thoracic crop of *L. japonicus* queens do not derive from, but would rather flow into, the gastral crop. As Hölldobler and Wilson (1990) suggested, the liquid stored in the thoracic crop might derive from converted queen tissues, such as metabolized fat bodies and flight muscles. Unmated, young queens of claustral species generally contain a far higher relative fat content than those of non-claustral species (Keller and Passera, 1989). Moreover, their fat content is depleted during claustral founding but not during mating flight (Passera et al., 1990; Tschinkel, 1993), suggesting that the fat reserves were utilized for the queen’s own survival, egg laying, and larval feeding during claustral ICF. However, in *Camponotus japonicus*, the founding queen would provide the first larvae with protein reserves rather than fat reserves (Idogawa et al., 2017). In addition, in queens of *Crematogaster opuntiae* (Myrmicinae) and *Camponotus festinatus*, protein reserves in both thoraces and gasters were also depleted during claustral ICF (Wheeler and Buck, 1996). The flight muscles were much enlarged in queens of claustral species than in those of non-claustral species (Keller et al., 2014), and thus, their thoraces with enlarged flight muscles were suggested to be a hallmark of specializations for claustral ICF (Peeters and Ito, 2015, 2001). The coordination between thoracic crop formation and flight muscle histolysis indicates that histolyzed flight muscles could provide a major source of thoracic crop content in *L. japonicus*. Further investigations are needed to reveal how such metabolized reserves are converted and stored in the thoracic crop.

We also found that during claustral ICF, some larvae cannibalized a sister egg (we could not discriminate between a reproductive or trophic egg) or larva. The ant larvae were suggested to be generally forced into a cannibalistic diet, especially oophagy, during claustral ICF (Urbani, 1991). Furthermore, the claustral queen of *S. invicta* and *Atta sexdens* (Myrmicinae) ingested trophic eggs, and then, the *Solenopsis* queen regurgitated it to larvae (Augustin et al., 2011; Cassill, 2002). Thus, the claustral queens would also convert a portion of their own metabolized body reserves into eggs, and then they may provide a portion of their eggs (or larvae) to existing larvae directly (Hölldobler and Wilson, 1990) or may regurgitate eggs ingested by themselves (Cassill, 2002; Meurville and LeBoeuf, 2021).

Our study described the process of thoracic crop formation in *L. japonicus* queens, which would be regulated to be spatiotemporally coordinated with flight muscle histolysis. Such developmental regulations for thoracic crop formation in a founding queen might be ubiquitous in claustral ICF species. In non-claustral ICF species, flight muscle histolysis occurs in founding queens (Haskins, 1941; Matte and Billen, 2021), but whether queen esophagus morphology changes during non-claustral ICF has not yet been examined. Conversely, thoracic crops have been reported in workers of various species, including non-claustral ponerine species (reviewed in Casadei-Ferreira et al., 2020). Therefore, developmental regulations for thoracic crop formation acquired in workers of more primitive ant lineages might have been co-opted in the founding queens of claustral ICF species. Future studies comparing founding queen internal structures, particularly those on the esophagus, during ICF between claustral and non-claustral species will enable us to examine the evolutionary process of thoracic crop in the claustral ICF ant queens.

## AUTHOR CONTRIBUTION

**Kurihara**: Investigation, Formal analysis, Visualization, Writing – original draft; **Ogawa**: Methodology, Investigation, Visualization, Writing – review & Editing; **Chiba**: Investigation, Writing – Review & Editing; **Hayashi**: Conceptualization, Funding acquisition, Writing – review & Editing; **Miyazaki**: Conceptualization, Funding acquisition, Project administration, Writing – review & Editing, Supervision.

## ACKNOWLEDGMENTS

We thank Hiromi Tanaka, Toshimi Hidaka, and other lab members in Tamagawa University for their assistance in field sampling, colony maintenance, and laboratory work. This work was supported by JSPS KAKENHI Grant Numbers JP19K06860 to SM and JP20K06816 to YH. The authors would like to thank Enago (www.enago.jp) for the English language review.

## FIGURE LEGENDS

**Figure S1**. Sample preparation process for TAFS microscopy. (A) A FAA-fixed sample was (B) decolorized, (C) transparentized, and then (D) sliced to 500 µm thicknesses.

**Figure S2**. Offspring number at each developmental stage (egg, larva, pupa, and adult worker) produced by *Lasius japonicus* founding queens during claustral ICF.

**Figure S3**. Larvae cannibalizing (A) egg and (B) larva in the claustral founding colonies. Scale bars 500 µm.

**Figure S4**. Changes in the surface structures of the thoracic esophagus in a decolorized sample. Dorsal view of the (A) queen thorax and dissected thoracic esophagus at (B, D) week 0 and (C, E) week 5 in claustral ICF.

## Notes

### Competing Interest Statement

The authors have declared no competing interest.

