## Supplementary figures and images for "Thoracic crop formation is spatiotemporally coordinated with flight muscle histolysis during claustral colony foundation of a *Lasius japonicus* queen"

### Fig. S1

#### Figure S1

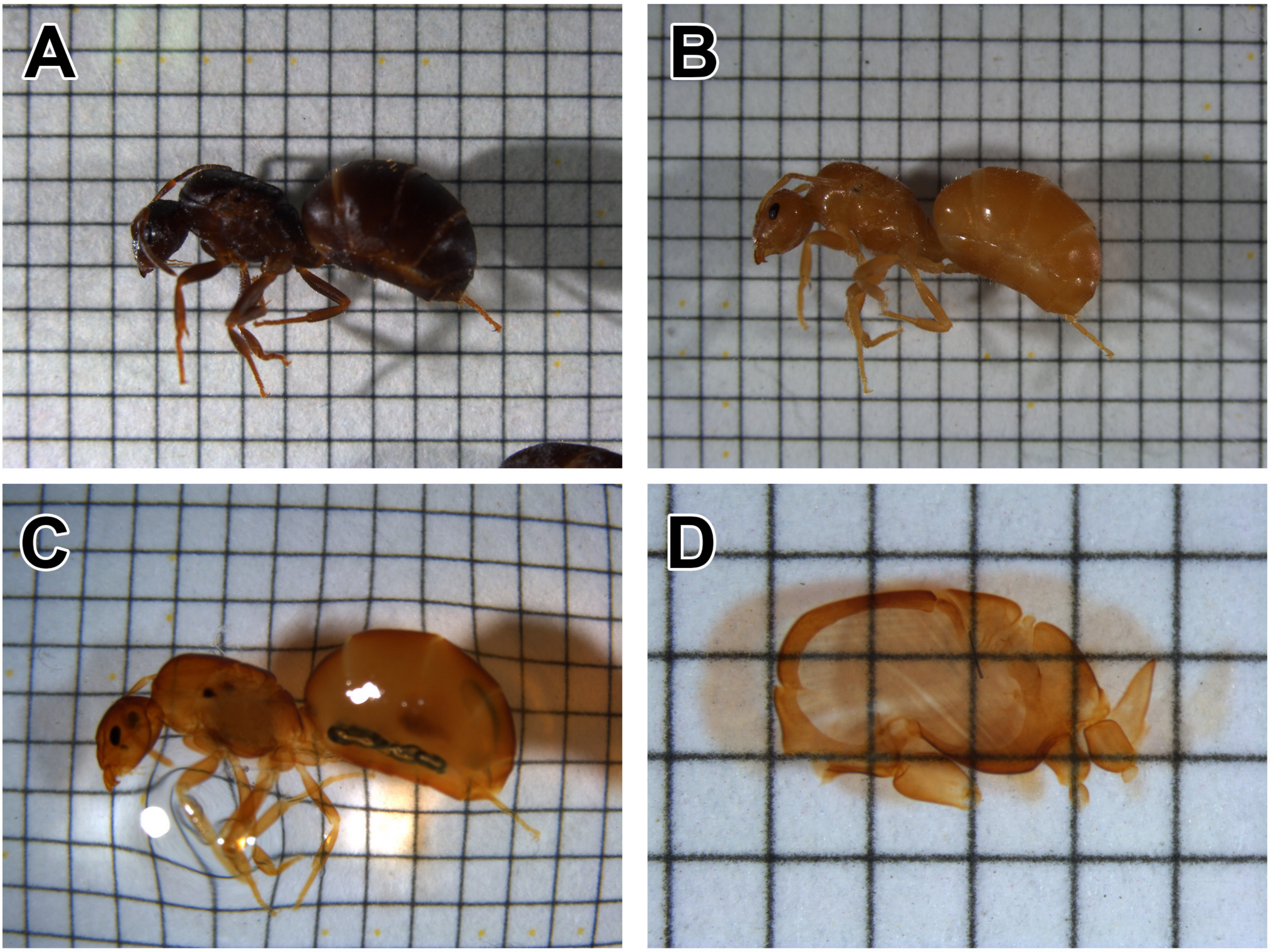

### Fig. S2

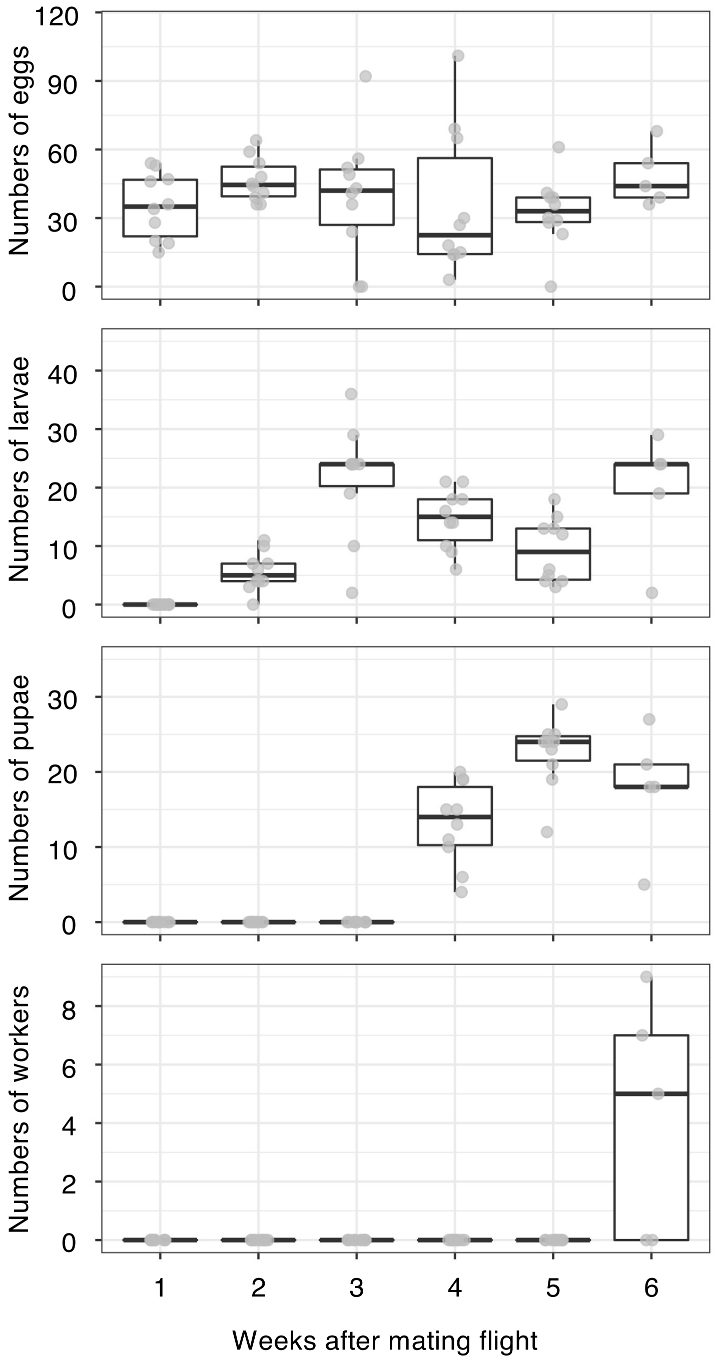

### Fig. S3

**Figure S3**

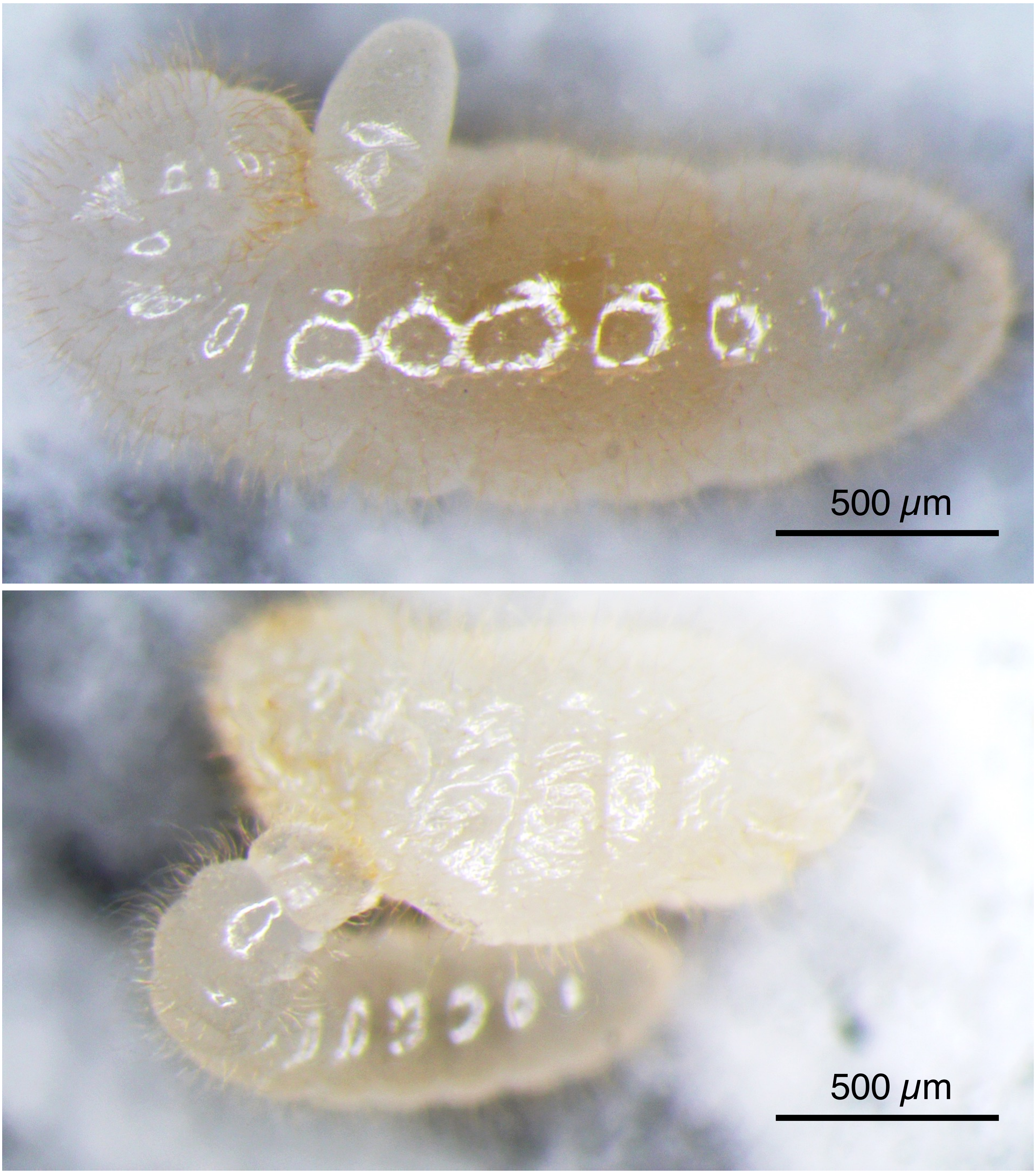

### Fig. S4

**Figure S4**

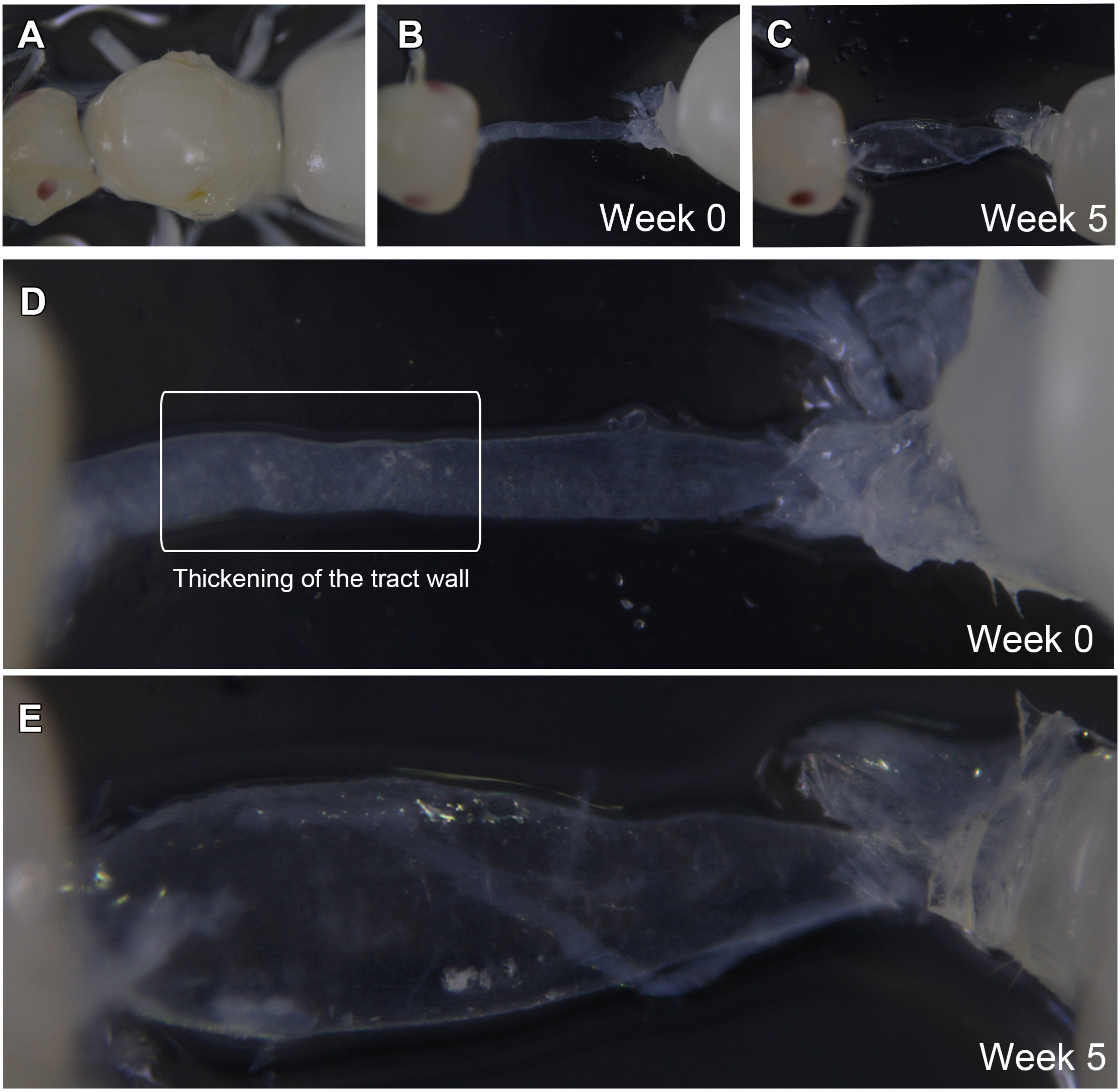
